# Convergent evolution of elaborate nests as structural defences in birds

**DOI:** 10.1101/2021.11.05.467427

**Authors:** Sally E. Street, Robert Jaques, Thilina N. De Silva

## Abstract

The ‘hanging-basket’ nests of some weaverbird and icterid species are among the most complex structures built by any animal, but why they have evolved remains to be explained. The precarious attachments and extended entrance tunnels characteristic of these nests are widely speculated to act as structural defences against invasion by nest predators, particularly tree-climbing snakes, but this hypothesis has yet to be systematically tested. We use phylogenetic comparative methods to investigate the role of nest structure in developmental period length, a proxy for offspring mortality, in the weaverbirds (Ploceidae) and icterids (Icteridae), two bird families in which ‘hanging-basket’ nests have independently evolved. We find that more elaborate nest designs, particularly those with entrance tunnels, are associated with longer developmental periods in both families. This finding is robust to potentially confounding effects of body mass, phylogenetic relationships and nest location. We provide the first comparative evidence for the role of elaborate nest designs in reducing offspring mortality from nest invasion. More broadly, our findings highlight the importance of offspring mortality risk in shaping diversity in birds’ nest designs and suggest that complex structures built by animals can buffer against environmental hazards and facilitate the evolution of slower life histories.

## Introduction

The structural complexity of birds’ nests varies enormously across species, from roughly constructed stick platforms to neatly-woven cups and domes (1–4). Why birds have evolved such diverse nest designs, however, remains poorly understood due to a surprising historical lack of research interest in the evolution of nest-building (5). The reasons for the evolution of even the most elaborate of all birds’ nests, the ‘hanging-baskets’ built by some species of weaverbirds (Ploceidae) and icterids (Icteridae), still remain to be established (1, 4). To build these nests, birds must knot, stitch and weave together hundreds of strips of nesting material (4, 6), requiring a significant amount of physical effort, manipulative skill and trial-and-error learning (6–9). Hanging-basket nests, therefore, must confer substantial fitness benefits to compensate for the costs of their construction (3). The primary advantage of hanging-basket nests is widely assumed to be protection from arboreal predators, particularly snakes (2, 4). Anecdotal evidence suggests that snakes struggle to access nests suspended from slim branches (9), and entrance tunnels may also prevent easy access by brood parasites (10, 11). While intuitive, this hypothesis is so far based largely on observational accounts and has yet to be investigated systematically in a phylogenetic comparative analysis.

Nest complexity varies considerably within the weaverbird and icterid families, making them ideal groups for testing hypotheses about the evolution of elaborate nest structures (**Figure 1**). In weaverbirds, nests range from roughly constructed, bulky masses firmly sited on thick branches to neatly-woven globes dangling precariously below slim vegetation, some with entrance tunnels up to a metre long (9, 12). In the icterids, nest complexity varies from typical songbird cups to suspended domes and elongated pendant-shaped nests, while some do not build nests at all, exploiting those built by other species instead (13). Despite their independent evolution, hanging-basket nests in the two families are highly similar in their design and construction: oropendolas and caciques use similarly intricate weaving methods, including some of the same stitches (e.g. half-hitches, loops and spiral binding) as some of the weaverbirds (4, 8). Such striking convergence strongly suggests common selection pressures; in both families, attacks by snakes and brood parasites are common and a significant source of offspring mortality (10–15).

**Figure 1.**
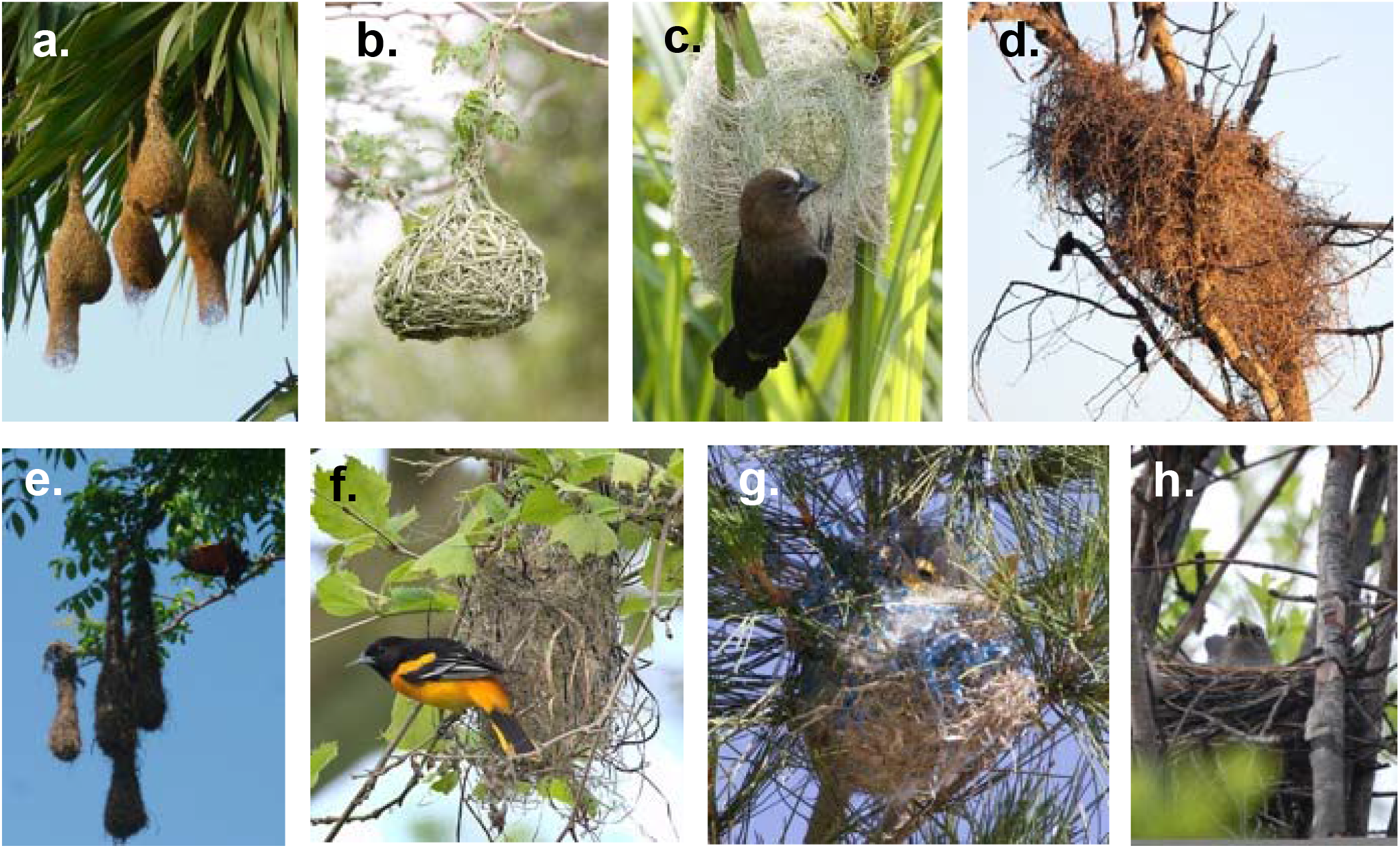
Examples illustrating the diversity of nest designs in weaverbirds (top row) and icterids (bottom row). a. Baya weaverbird (Ploceus philippinus), b. Southern masked weaver (Ploceus velatus) c. Thick-billed weaver (Amblyospiza albifrons), d. Red-billed buffalo weaver (Bubalornis niger); e. Montezuma oropendola (Psarocolius Montezuma), f. Baltimore oriole (Icterus galbula), g. Bullock’s oriole (Icterus bullocki), h. Rusty blackbird (Euphagus carolinus). Image credits: a. Pinakpani - CC BY-SA 4.0, https://commons.wikimedia.org/w/index.php?curid=59438503 b. Chris Eason – CC BY 2.0, https://commons.wikimedia.org/w/index.php?curid=4127816 c. Derek Keats – CC BY 2.0, https://commons.wikimedia.org/w/index.php?curid=45087290 d. Derek Keats – CC BY 2.0, https://www.flickr.com/photos/dkeats/6041784294 e. Caspar S - CC BY 2.0, https://commons.wikimedia.org/w/index.php?curid=41959845 f. Andrew Weitzel - CC BY-SA 2.0, https://commons.wikimedia.org/w/index.php?curid=105254878 g. HarmonyonPlanetEarth – CC BY 2.0, https://commons.wikimedia.org/w/index.php?curid=45578683 h. Robin Corcoran, USFWS – CC0, https://pixnio.com/fauna-animals/birds/blackbirds-pictures/female-rusty-blackbird-euphagus-carolinus-on-nest. All images have been edited only by cropping and re-sizing.

Here, we conduct the first systematic test of the hypothesis that elaborate ‘hanging-basket’ nests in birds have evolved as structural defences, using phylogenetic comparative methods. If elaborate structural features of hanging-basket nests provide protection from nest invasion by arboreal predators or brood parasites, then species building more protected nests, i.e. those with tunnels and/or with more precarious attachments, should show evidence of reduced offspring mortality compared with species that build less protected nests. The evolution of species’ life histories is strongly shaped by extrinsic mortality risks (16), and therefore life histories can be used as proxies for evolutionary responses to predation in comparative analyses (as in e.g. (17, 18)). The length of time that developing offspring spend in the nest is of particular relevance to the present study. Theoretical analyses suggest that selection should favour rapid maturation where offspring are raised in exposed locations, while offspring raised in protected nests can afford to develop more slowly due to relaxed predation pressure (19). In support of this model, previous comparative analyses have shown that nestling period length correlates negatively with offspring mortality from nest predation (20) and brood parasitism (21). We test predictions by examining the effects of nest design on species’ developmental period length (incubation, nestling and their combined duration), accounting for potentially confounding effects of phylogeny, body mass and nest location.

## Methods

### Data collection

We obtained data on nest design, life history traits and body mass in weaverbird and icterid species from multiple secondary sources. We classified variation in nest design primarily based on descriptions and images of nests from Birds of the World Online (12, 13). We also obtained photographs of nests from the Natural History Museum at Tring, both via the NHM Data Portal (22) and from an in-person visit. After finding that information on weaverbird nests in Birds of the World Online was less comprehensive than for icterids, we consulted two additional sources for descriptions and images of weaverbird nests: PHOWN, a citizen science project collating photographs of weaverbird nests (23) and a comparative study of weaverbird nests conducted prior to the development of modern phylogenetic methods (9). Where information conflicted between sources, we generally prioritised photographs over textual descriptions, taking into account image quality. Where there was insufficient information to classify species’ nest designs (i.e. vague textual descriptions, poor quality images or no information at all), we conducted further targeted Google Image and Google Scholar searches (using the search string “[binomial] OR [common name] AND nest”) for further information. If we could find no reliable further information from these targeted searches, we excluded species from the analyses.

We classified weaverbird nests based on two separate design features: the presence of entrance tunnels, and the type of attachment (**SI Figure 1**). We considered nests to have tunnels if an external, tube-shaped extended entrance was clearly present, of any length. We did not count short extensions to the upper side of the nest entrance (often referred to as ‘porches’ or ‘lips’) as tunnels. We treated nest attachment as a categorical variable with three levels: ‘supported’, ‘suspended’ or ‘pendulous’, increasing in precariousness and presumed difficulty of access by predators. We classified as ‘supported’ nests those that are attached from the underside to a branch, built on the ground or firmly attached on two sides between vertical supports rising up from the ground. We classified as ‘suspended’ nests attached at the top or side(s), so that the bulk of the nest lies below the substrate, while we classified ‘pendulous’ nests as those hanging from the substrate above by a single point of attachment. For icterids, we treated nest design as a single three-level factor, ordered by precariousness of attachment (supported < suspended < pendulous, **SI Figure 1**). We classified as ‘supported’ icterid nests that are firmly attached to vegetation or other substrates from below (including nests built on the ground and inside cavities). We classified as ‘suspended’ bag-shaped pouches attached at the rim or by multiple ‘straps’, such that the bulk of the nest hangs below the substrate, while more elongated, pendant-shaped nests as ‘pendulous’. ‘Pendulous’ nests in icterids have entrances at the top rather than the bottom as in weaverbirds (**Figure 1e, SI Figure 1**), creating an upward-facing entrance tunnel. Therefore, we could not classify tunnels as a design feature distinct from attachment type in the icterids as we did for the weaverbirds. We excluded icterid species that do not build their own nests, including brood-parasitic cowbirds (n=5) and troupials reliant on the nests of other species (n=2). Where there was intraspecific variation in nest design, we considered a feature to be present if it occurs within the typical range of nest designs for that species. For example, we considered tunnels present in a species that builds nests both with and without tunnels, and pendulous nests present in a species that builds both pendulous and suspended nests. Where available, we also recorded the maximum recorded entrance tunnel length for tunnel-building weaverbirds, and maximum recorded nest length (both in cm) in suspended or pendulous nest-building icterids.

Nest location may play at least as important a role as nest structure in protecting developing offspring from potential nest invaders (3). Weaverbirds and icterids often build their nests in locations inaccessible to most terrestrial predators, attached precariously to the tips of slim branches high off the ground (9, 12, 13). Building nests in thorny vegetation, over water, in large breeding colonies and/or close to the nests of aggressive stinging insects or predatory birds may also provide effective deterrents to a wide range of potential predators, regardless of nest structure (9). An observational study of baya weaverbirds in India found that characteristics of nest location were in fact more strongly associated with fledging success than were those of nest structure (10). Therefore, it was important to consider both aspects of nest structure and location in our analyses, particularly as they may be confounded with one another if elaborate nests tend to be built in more protected locations. Along with nest structure data, we collected data on potentially protective features of nest location including nest height, colonial nesting, nesting in thorny vegetation, nesting over water, nesting in association with raptors and nesting in association with stinging insects from Birds of the World Online (12, 13). Nest height was given in metres above the ground or water. Where ranges or multiple values were provided for nest height, we took the median of the minimum and maximum provided, otherwise we used single values. We treated ground-nesting as nesting at a height of zero metres. We counted as colonial nesters species that breed in colonies, regardless of colony size, and including those described as “loosely” or “semi” colonial, as well as those that nest both in colonies and solitarily. We did not count as colonial, however, species with uncertain descriptions of coloniality such as “appears to be colonial” or “presumably colonial”. We classified species as nesting in thorny vegetation, over water, in association with stinging insects (e.g. ants, wasps, bees or hornets) and in association with raptors if there was at least one clear description or image of each relevant behaviour. We could not analyse the potential role of nesting associations with raptors in developmental period length as it was reported in a very small number of species in our samples (n=6 in weaverbirds, none in icterids). Additionally, we found an insufficient number of icterid species nesting in association with stinging insects (n=3) to allow for statistical analysis.

We obtained data on incubation period duration (days), nestling period duration (days) and adult body mass (grams) primarily from Birds of the World Online (12, 13). We used female body mass where available, otherwise we used male body mass or body mass of unknown sex. We included data from captive populations to increase sample size, though selected estimates from the wild when both data from wild and captive populations were available. Where life history traits were reported as ranges or multiple values, we took the median of the minimum and maximum estimates, otherwise we used single values. We resolved mis-matches between species’ names in different datasets and the phylogenies where possible by referring to the latest Bird Life International taxonomy (24). Because we found that we had nest data for more species than we had life history data, we performed further targeted literature searches for additional life history data to increase sample sizes. Here, we checked two large comparative avian life history databases (25, 26) and performed a Google Scholar search for each individual species’ binomial and common names with relevant life history keywords (using the search string “[binomial] OR [common name] AND incubation OR nestling OR fledging”). As a result, we obtained life history data for an additional 6 weaverbird and 7 icterid species. A complete list of sources of additional life history data is available in **SI Table 1**.

### Data analysis

To investigate whether developmental periods are longer in species building more protected nests, we fit regression models in which either incubation period length, nestling period length or total developmental period length (summed incubation and nestling periods) was the outcome variable, with nesting variables as predictors. We also included body mass as a predictor in all models to control for allometric scaling of developmental periods with body size (26). We use p-values to estimate the probability of the observed effects under the null hypotheses of regression coefficients of zero. We do not, however, specify any arbitrary thresholds for ‘statistical significance’ in advance since p-values are continuous quantities (27). We log-10 transformed all continuous variables prior to analysis as they were generally positively skewed. For all analyses, we examined standard regression diagnostic plots and found no concerning patterns.

We used phylogenetic generalised least squares (PGLS) regression to account for the non-independence of species data caused by phylogenetic relationships (28, 29). PGLS analyses adjust regression coefficients according to the degree of phylogenetic influence in model residuals, estimating phylogenetic signal using Pagel’s λ (28, 29). λ varies from 0 to 1, where 0 indicates no phylogenetic signal and 1 the maximum possible signal, assuming a Brownian motion model in which the amount of phenotypic change is directly proportional to evolutionary time (30, 31). We obtained a near-species-level multi-locus phylogeny for the weaverbirds based on 4 mitochondrial markers and 4 nuclear introns (32, 33), constructed using Bayesian inference in BEAST v.1.82 (34), executed with BEAGLE (35) and summarised the tree block in the form of a maximum-clade credibility tree. For icterids, we used a complete phylogeny based on 2 mitochondrial genes, 4 nuclear genes and whole mitochondrial genomes for selected species, constructed using maximum likelihood estimation (36).

Most of the weaverbird analyses reported λ values of 0, which was unexpected given that developmental periods are generally strongly influenced by phylogeny in birds (26). Low λ values have a range of possible explanations including both biological factors, such as highly labile traits (37) and statistical artefacts, particularly low power (30). In our case, the latter was unlikely as we do find >0 values of λ in some weaverbird models (and in models based on smaller samples of icterids), and λ values are still 0 or relatively low (≤0.50) even when estimating phylogenetic signal in each life history trait individually. We investigated this issue further by estimating Pagel’s δ, which allows the speed of evolutionary change to vary through time, along with Pagel’s λ (31). δ varies from 0 to 3, where values of <1 suggest faster evolutionary change earlier in the phylogeny, consistent with adaptive radiations, while higher values suggest faster change later in the phylogeny, suggestive of recent convergent adaptations (31). We found high (>2.2) values of δ for all three life history traits in weaverbirds. We therefore concluded that the low λ values for developmental periods in weaverbirds are most likely indicative of unusually labile developmental periods in this clade, consistent with recent, rapid convergent evolution. This interpretation is supported by substantial disagreements recently identified between morphological and phylogenetic affinities in the weaverbirds (32).

For illustrative purposes, we also performed ancestral states reconstructions on nest tunnels in weaverbirds and nest attachment in icterids. We used maximum likelihood methods to estimate transition rates, fitting a simple model in which rates of gains and losses were assumed to be equal (38). We ran all analyses in R (39), using functions from the *caper* (40), *ape* (38) *phangorn* (41) and *phytools* (42) packages. After matching species between the datasets and phylogenies, 56 weaverbird and 48 icterid species remained with complete data on nest design and developmental periods. Of these, 15 weaverbird and 17 icterid species also had data on tunnel or nest length respectively. All data and code are accessible from DataDryad at the following link: https://datadryad.org/stash/share/YRYnZIGry7E9kebGYFwEbNp2UADJRPsh2LxIUuWf7JE.

## Results

**Figure 2** illustrates variation in nest design across weaverbird and icterid species and displays the results of ancestral states reconstructions.

**Figure 2.**
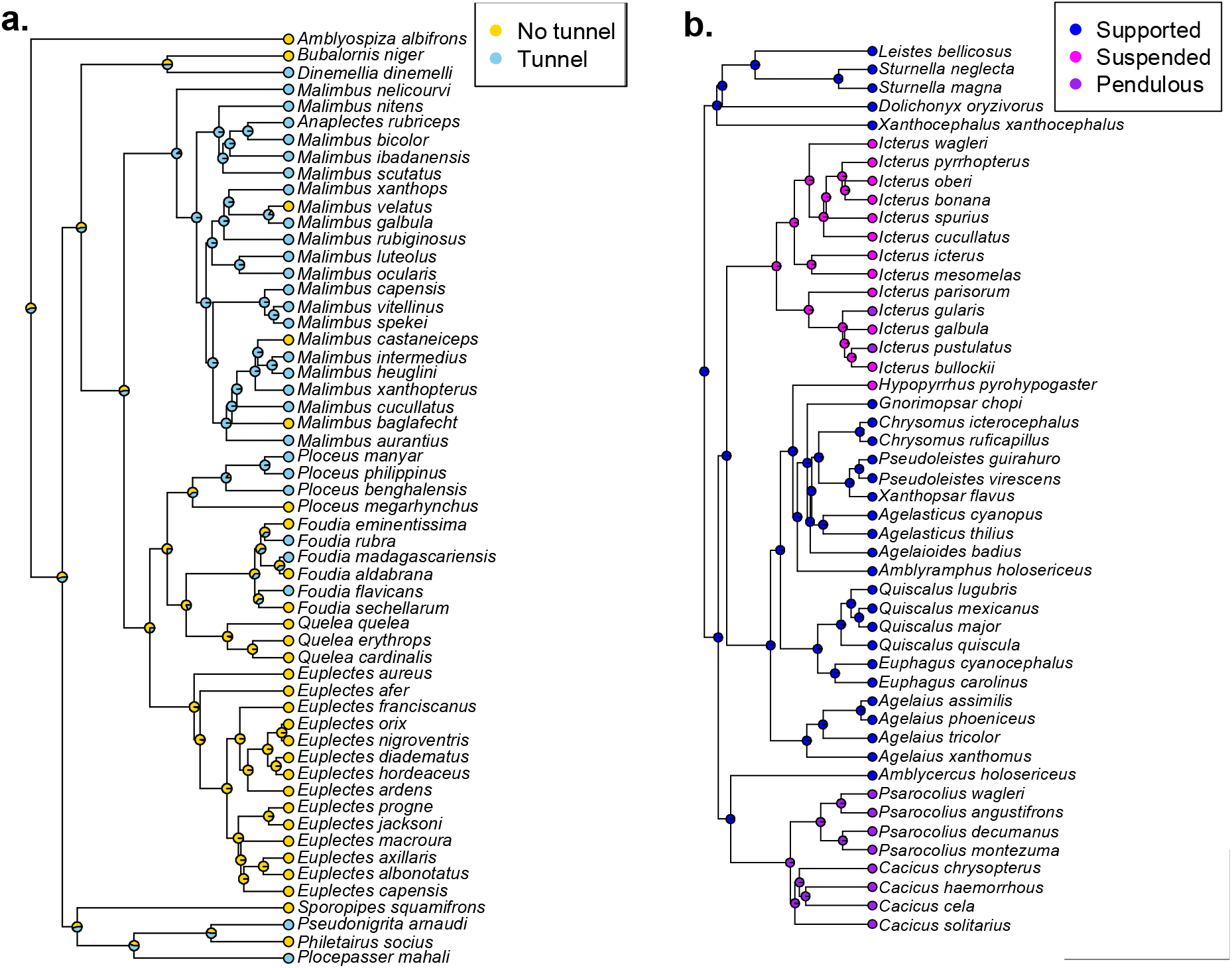
Nest characteristics mapped onto phylogenies for a. weaverbirds and b. icterids. Tip labels show observed nest classifications among extant species, while node labels illustrate the results of the ancestral states reconstruction, with shaded areas of the pie charts indicating the probability of each state at each node.

### Weaverbirds

Weaverbird species building nests with entrance tunnels have offspring with slightly longer combined developmental periods than those building nests without entrance tunnels (**Figure 3a, Table 1**). Predicted values based on model coefficients suggest that building a nest with a tunnel is associated with an additional ~1.3 days from laying to fledging age, for a weaverbird species of average body mass. When separating developmental periods into incubation and nestling periods, we find that tunnels are associated with relatively longer incubation periods rather than nestling periods (**Table 1**). Developmental periods, however, do not differ between weaverbird species building pendulous, suspended or supported nests (**Table 2**). Tunnel length and developmental period length appear not to be positively correlated, although statistical power is low due to the small sample size (**Figure 3c, Table 3**). Nest height is not related to developmental period length (**SI Table 2**), and nesting in protected locations is generally not associated with longer developmental periods (**SI Tables 3-7**), apart from longer incubation periods in species nesting over water (**SI Table 5**). Positive, though weaker, effects of both tunnels and nesting over water remain when both are included as predictors in the same model (**SI Table 6**).

**Table 1.** Full results from models predicting log10 developmental period length, log10 incubation period length and log10 nestling period length from tunnel presence and log10 body mass in weaverbirds.

| Dep. variable | Ind. variable | Estimate | S.E. | T-value | p-value | N | R<sup>2</sup> | λ |
| --- | --- | --- | --- | --- | --- | --- | --- | --- |
| Dev. period | Tunnel | 0.02 | 0.01 | 1.60 | 0.11 | 56 | 0.18 | 0.00 |
|  | Body mass | 0.10 | 0.04 | 2.61 | 0.01 |  |  |  |
| Incubation period | Tunnel | 0.02 | 0.01 | 2.14 | 0.04 | 58 | 0.12 | 0.00 |
|  | Body mass | 0.04 | 0.03 | 1.31 | 0.20 |  |  |  |
| Nestling period | Tunnel | 0.02 | 0.02 | 0.94 | 0.35 | 57 | 0.13 | 0.00 |
|  | Body mass | 0.14 | 0.06 | 2.43 | 0.02 |  |  |  |

**Table 2.**
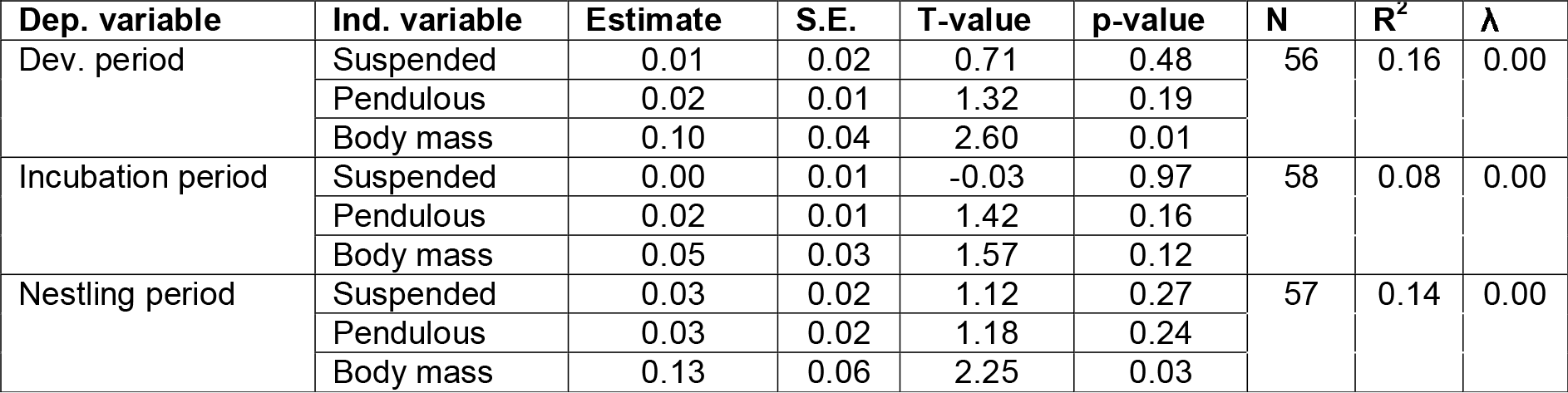
Full results from models predicting log10 developmental period length, log10 incubation period length and log10 nestling period length from attachment type and log10 body mass in weaverbirds.

| Dep. variable | Ind. variable | Estimate | S.E. | T-value | p-value | N | R<sup>2</sup> | λ |
| --- | --- | --- | --- | --- | --- | --- | --- | --- |
| Dev. period | Suspended | 0.01 | 0.02 | 0.71 | 0.48 | 56 | 0.16 | 0.00 |
|  | Pendulous | 0.02 | 0.01 | 1.32 | 0.19 |  |  |  |
|  | Body mass | 0.10 | 0.04 | 2.60 | 0.01 |  |  |  |
| Incubation period | Suspended | 0.00 | 0.01 | -0.03 | 0.97 | 58 | 0.08 | 0.00 |
|  | Pendulous | 0.02 | 0.01 | 1.42 | 0.16 |  |  |  |
|  | Body mass | 0.05 | 0.03 | 1.57 | 0.12 |  |  |  |
| Nestling period | Suspended | 0.03 | 0.02 | 1.12 | 0.27 | 57 | 0.14 | 0.00 |
|  | Pendulous | 0.03 | 0.02 | 1.18 | 0.24 |  |  |  |
|  | Body mass | 0.13 | 0.06 | 2.25 | 0.03 |  |  |  |

**Table 3.** Full results from models predicting log10 developmental period length, log10 incubation period length and log10 nestling period length from log10 nest tunnel length and log10 body mass in weaverbirds.

| Dep. variable | Ind. variable | Estimate | S.E. | T-value | p-value | N | R<sup>2</sup> | λ |
| --- | --- | --- | --- | --- | --- | --- | --- | --- |
| Dev. period | Tunnel length | 0.03 | 0.03 | 0.86 | 0.41 | 15 | 0.06 | 0.00 |
|  | Body mass | -0.01 | 0.12 | -0.09 | 0.93 |  |  |  |
| Incubation period | Tunnel length | 0.02 | 0.02 | 0.98 | 0.35 | 15 | 0.19 | 0.00 |
|  | Body mass | 0.10 | 0.08 | 1.22 | 0.25 |  |  |  |
| Nestling period | Tunnel length | 0.03 | 0.05 | 0.62 | 0.55 | 15 | 0.05 | 0.00 |
|  | Body mass | -0.10 | 0.17 | -0.59 | 0.56 |  |  |  |

**Figure 3.**
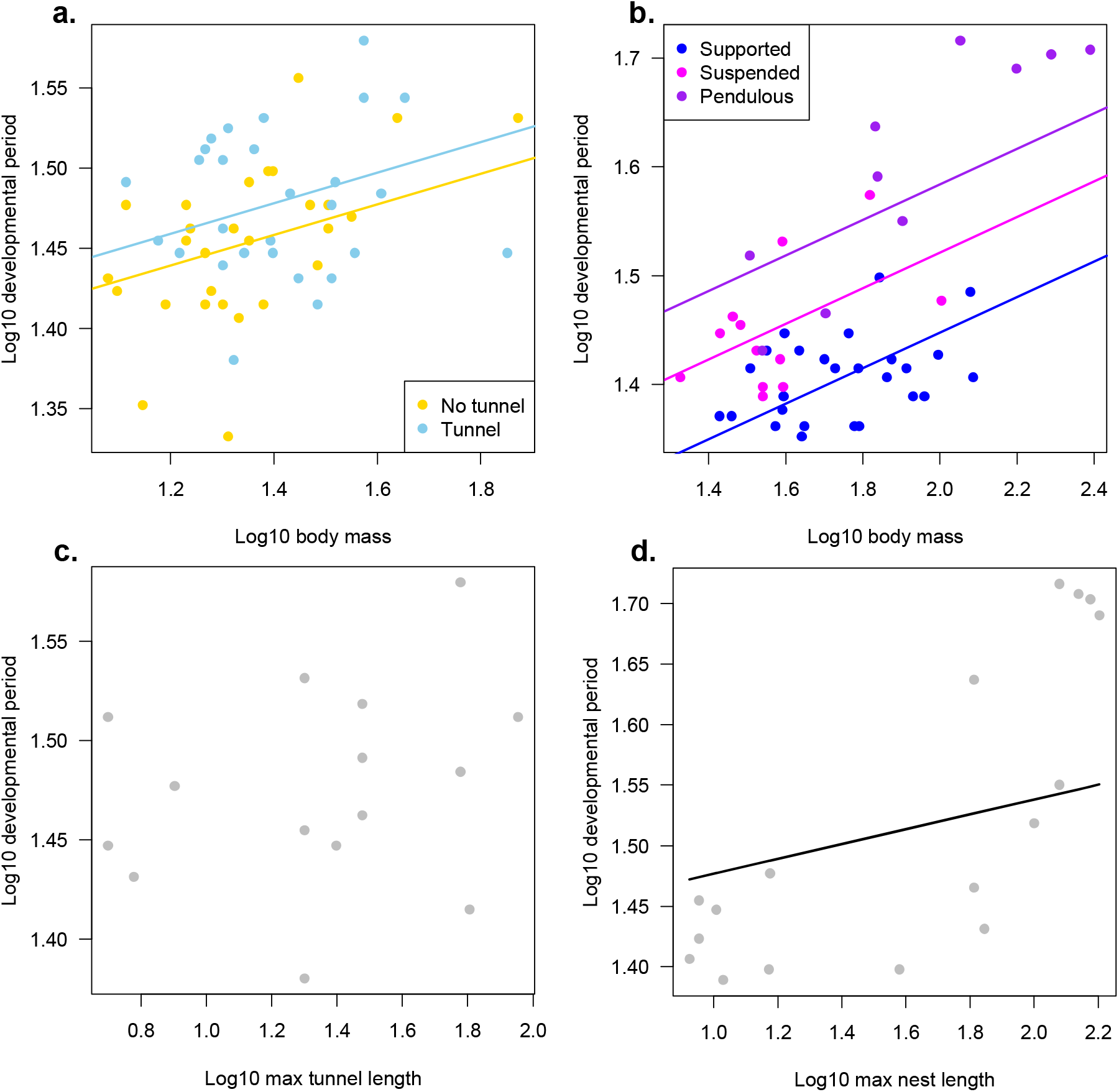
Scatterplots illustrating a. longer developmental periods relative to body mass in weaverbird species building nests with tunnels than those building nests without tunnels b. increasing developmental periods relative to body mass in icterids building supported, suspended and pendulous nests, c. the relationship between developmental period length and tunnel length in weaverbird species and d. a positive association between developmental period length and nest length in icterid species. Fit lines are plotted using PGLS model coefficients. The fit line on panel d. is calculated assuming an icterid species of average body mass.

### Icterids

Icterid species building pendulous nests (which incorporate upward-facing tunnels) have longer developmental periods compared with those building suspended nests, who in turn have relatively longer developmental periods than those building supported nests (**Figure 3b, Table 4**). Model predictions suggest that the offspring of pendulous nest-building species require an additional ~4.7 days to reach fledging age compared with suspended nest-builders, who in turn take ~4.7 more days than supported nest builders, assuming an icterid species of average body mass. In contrast to the weaverbirds, nest type has a stronger effect on the length of nestling than incubation periods (**Table 4**). Nest length is positively, although fairly weakly, correlated with developmental period length, particularly nestling period, controlling for body mass (**Table 5**). Developmental period length (both incubation and nestling periods) also increases with nest height off the ground in icterids (**SI Table 8**). However, effects of nest type on developmental period length remain when both nest type and nest height are included in the same model (**SI Table 9**). Similarly to weaverbirds, in icterids we find no evidence that nesting in other protected locations is associated with longer developmental periods (**SI Table 10-12**).

**Table 4.**
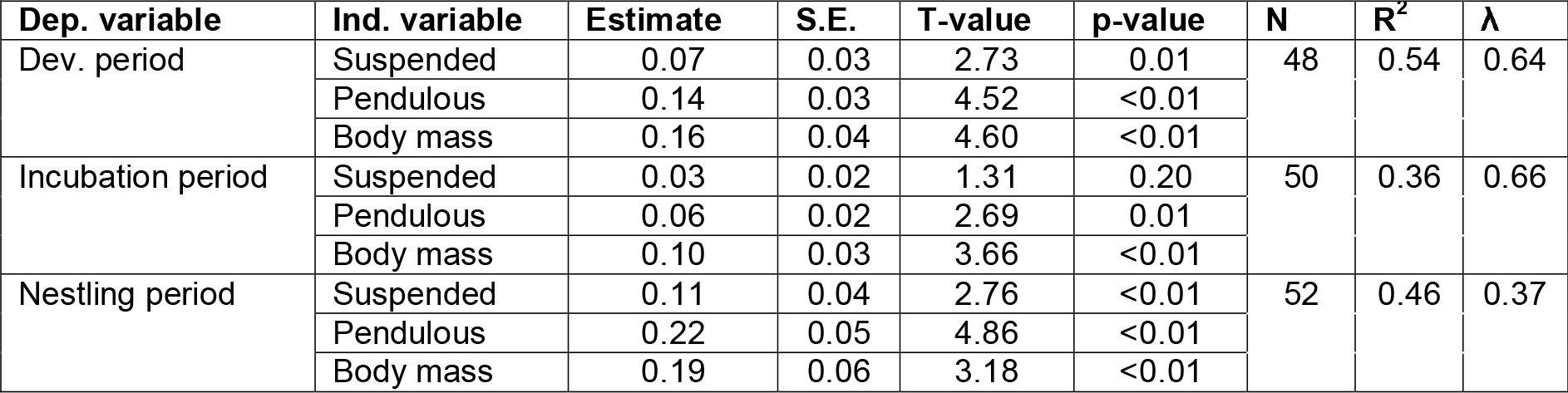
Full results from models predicting log10 developmental period length, log10 incubation period length and log10 nestling period length from nest type and log10 body mass in icterids.

**Table 5.**
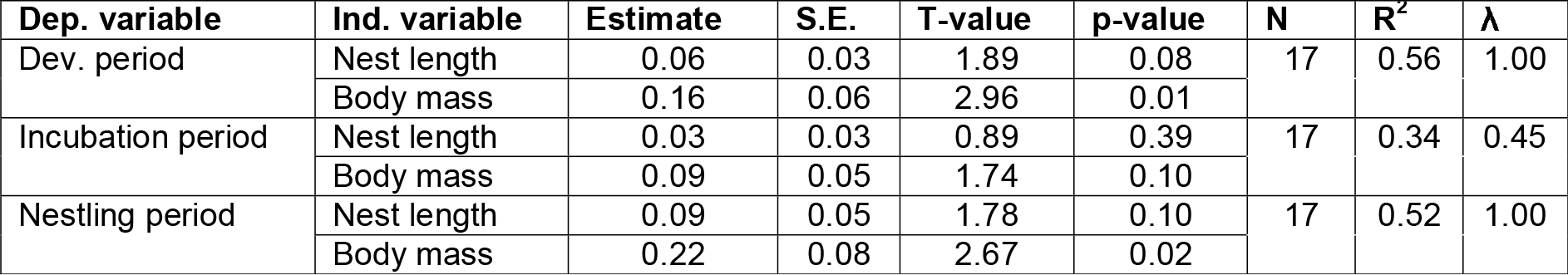
Full results from models predicting log10 developmental period length, log10 incubation period length and log10 nestling period length from log10 nest length and log10 body mass in icterids.

| Dep. variable | Ind. variable | Estimate | S.E. | T-value | p-value | N | R<sup>2</sup> | λ |
| --- | --- | --- | --- | --- | --- | --- | --- | --- |
| Dev. period | Nest length | 0.06 | 0.03 | 1.89 | 0.08 | 17 | 0.56 | 1.00 |
|  | Body mass | 0.16 | 0.06 | 2.96 | 0.01 |  |  |  |
| Incubation period | Nest length | 0.03 | 0.03 | 0.89 | 0.39 | 17 | 0.34 | 0.45 |
|  | Body mass | 0.09 | 0.05 | 1.74 | 0.10 |  |  |  |
| Nestling period | Nest length | 0.09 | 0.05 | 1.78 | 0.10 | 17 | 0.52 | 1.00 |
|  | Body mass | 0.22 | 0.08 | 2.67 | 0.02 |  |  |  |

## Discussion

We find that in both weaverbirds and icterids, species building more elaborate nests, particularly those with extended entrance tunnels, have offspring with longer developmental periods. Since theoretical and comparative evidence shows that offspring develop more slowly when offspring mortality is lower (16, 19–21), these results are consistent with the hypothesis that nests with extended entrance tunnels limit the exposure of developing broods to nest invaders. The consistency of these findings is striking given that ‘hanging-basket’ nests have evolved independently in the weaverbirds and icterids. We also find that in icterids at least, developmental period length is positively correlated with nest tunnel length, suggesting that longer tunnels are more effective at hindering access by nest invaders. We find some evidence that nesting in protected locations is also associated with longer developmental periods in these two families, including nesting over water in weaverbirds and nesting higher off the ground in icterids. However, none of the effects of nest structure on developmental periods is confounded by nest location. Therefore, we provide the first comparative evidence in favour of the long-held hypothesis that elaborate hanging-basket nests in birds have evolved as structural defences against nest invasion.

While our results do not directly demonstrate which specific nest invaders elaborate nests have evolved in response to, they do suggest that brood parasites may play a more important role than previously appreciated. Both precarious attachments and extended entrance tunnels should protect against attacks by arboreal snakes, yet in weaverbirds, where the two features are separable, we find that longer developmental periods are associated only with tunnels, not attachment type. Tunnels should make it physically more difficult for brood parasites to access nests quickly and avoid detection by hosts (10, 11), while precarious nest attachments are of no obvious relevance to ease of access by brood parasites. The effect of tunnels but not attachment type on developmental periods in weaverbirds, therefore, is more consistent with protection against brood parasites than arboreal snakes. The finding that nest tunnels are associated with longer incubation periods rather than nestling periods in the weaverbirds is also consistent with an important role of nest tunnels in protection from brood parasites. A previous comparative analysis has shown that egg colouration is less variable in tunnel-building than non-tunnel-building *Ploceus* weavers, consistent with relaxed parasite pressure on species building structural defences (11). Further, pendulous nests in the icterids are not obviously well-designed to prevent access by snakes since the nest entrance is at the top, allowing for relatively easy access from branches above. The potential role of brood parasitism in elaborate nest designs has so far been overlooked in comparison to snake predation, but remains a plausible explanation given the high risk of parasitism in many weaverbird and icterid species. The diederik cuckoo (*Chrysococcyx caprius*) alone targets at least 34 different host weaverbird species (43), with rates of parasitism as high as 50% in some populations (e.g. southern red bishops, *Euplectes orix* (14)). The vast majority of icterids are parasitized by at least one cowbird species (44), with rates of close to 100% parasitism not infrequently reported (e.g. orchard orioles, *Icterus spurius* (15)). Brood parasitism, therefore, likely exerts significant selection pressures on nest design in weaverbird and icterids.

In contrast to nest structure, we find that nest location has generally little effect on offspring developmental periods across weaverbird and icterid species, other than nesting over water in weaverbirds and nesting high off the ground in icterids. This result is perhaps surprising given that many prior observational studies have found that multiple aspects of nest location, particularly nest height, affect exposure to predators (3). Our results appear to conflict, for example, with a prior study of baya weaverbirds which found that both nesting at greater heights and in thorny trees increased the probability of fledging success, while entrance tube length had no significant effect (10). This study, however, seemed to capture only a limited amount of variation in entrance tube length among baya weavers: nests within the study had entrance tubes up to 14cm long while they can reach as long as 90cm in this species (10). Baya weavers, further, may deviate from general patterns in the weaverbirds as they are not parasitized by cuckoos and may be affected more by rodent than snake predation (10). A greater number of population-level studies on the role of nest structure and location in offspring survival across a wider diversity of weaverbird species would therefore be valuable for a more detailed understanding of how varying levels of predation and brood parasitism affect the design of elaborate nests both within and across species. Our findings also seem to conflict with those of a recent global-scale comparative analysis in birds which found that developmental durations increase with nest height, but not with more protected nest structures (26). This study, however, focused on broad-scale comparisons between open and closed nest structures rather than on more elaborate features, such as tunnels, which are found only in a limited number of bird families. The weaverbirds and icterids likely deviate from these general patterns due to the exceptionally elaborate designs found in these families.

Birds’ nest design is undoubtedly influenced by multiple selection pressures in addition to predation and brood parasitism, particularly climatic conditions (1–3). However, while protection from heat, wind and rain could conceivably help explain why weaverbirds and icterids nests are built from strong, tightly woven fabric (9), climate has no obvious connection to the construction of entrance tunnels. Elaborate nest designs in birds may also be shaped by mate choice in some species (3), a suggestion that appears compelling for weaverbirds given that unusually for birds, the most elaborate nests are typically built solely by males (9, 12). In some species, such as Village weavers (*Ploceus cucullatus*), males appear to draw attention to the quality of their constructions by hanging upside down from nests during mating displays, and females seem to carefully inspect the neatness of their weaving before accepting the nest (45). However, similar sexual selection pressures are unlikely to explain the convergence of elaborate nests in the weaverbirds and icterids since hanging-basket nests are predominantly built by females rather than males in the latter (4, 13). The idea that sexual selection has shaped the design of hanging-basket nests is yet to be investigated in a comparative analysis, although population-level studies have so far not supported it. In baya weavers, female choice is influenced primarily by nest location rather than nest structure (46), and in Village weavers, while females prefer nests made with fresher material, their choices are primarily determined by male behaviour and mate quality (45, 47). Overall, therefore, while climate and sexual selection are generally potentially important drivers of nest design in birds, protection from predators and/or brood parasites appears the most plausible explanation for the evolution of elaborate nest structures in the weaverbirds and icterids.

We find evidence that hanging-basket nests, which are among the most elaborate structures built by any animal, have evolved convergently in two bird families in response to similar threats to offspring survival. Ours is the first comparative study to identify potential causes of the evolution of such elaborate nests since John Crook’s foundational study in the 1960s (9), taking advantage of the development of modern phylogenetic comparative methods and increasing availability of molecular phylogenies. We therefore confirm a long-held, but until now untested, assumption that entrance tunnels in elaborate birds’ nests function as structural defences against nest invasion. More broadly, our findings support the idea that by constructing protective structures, some animal species can exert greater control over their exposure to environmental hazards through behaviour, lowering offspring mortality and facilitating the evolution of slower life histories (48, 49). The ability to construct protective environments may have similarly contributed to extended life histories in several other independently evolved animal lineages, such as eusocial mammals and insects, and even humans.

## Author contributions

SES designed the study, collected data, analysed data and wrote the manuscript. RJ obtained preliminary datasets, conducted preliminary analysis, contributed to study design and contributed to writing the manuscript. TNDS provided and advised on the use of the weaverbird phylogeny, contributed to study design and writing the manuscript.

## Acknowledgements

We are grateful to Dr Alexis Powell for providing the icterid phylogeny, Mr Douglas Russell at the Natural History Museum at Tring for providing access to the nest collection and the Association for the Study of Animal Behaviour for funding the study through a Research Grant to SES.

