## Supplementary Information for "Convergent evolution of elaborate nests as structural defences in birds"

**SI FIGURES**

**SI Figure 1**

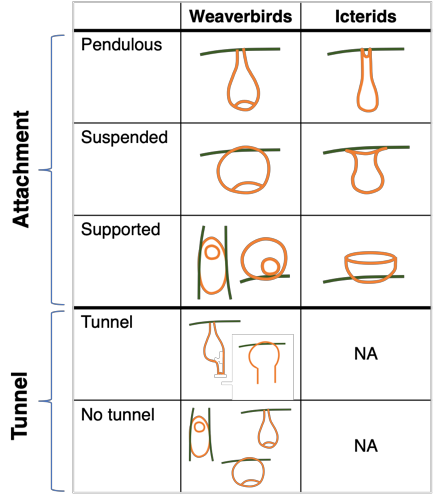

*Diagram illustrating the classification scheme used for nest design in the weaverbirds (both attachment and tunnels coded separately) and icterids (attachment and tunnel combined into a single nest design variable – pendulous nests incorporate upward facing tunnels in the icterids).*

**SI TABLES**

**SI Table 1**

| **Family** | **Species** | **Data** | **Source** |
| --- | --- | --- | --- |
| Weaverbirds | *Dinemellia dinemelli* | Incubation | Bagley-Vyas (2014), Avian Scientific Advisory Group Species Fact Sheet, <http://aviansag.org/Fact_Sheets/PACCT/White-Headed_Buffalo_Weaver.pdf> |
| Weaverbirds | *Foudia eminentissima* | Incubation, nestling & body mass | Cooney et al. (2020). Ecology and allometry predict the evolution of avian developmental durations. *Nature Communications*, 11 (2383), <http://doi.org/10.1038/s41467-020-16257-x> |
| Weaverbirds | *Foudia sechellarum* | Nestling | Vega et al. (2007), Extreme gender-based post-fledging brood division in the toc-toc, *Behavioral Ecology*, 18: 730-735,  <https://doi.org/10.1093/beheco/arm038> |
| Weaverbirds | *Malimbus scutatus* | Incubation & nestling | Egwumah & Faga (2013), Effects of early and late-dry season fires on mortality, dispersal and breeding of malimbe *Malimbus scutatus* in the Southern Guinea savannah, *International Journal of Biological and Chemical Sciences*, 6: 3082-3088, <https://www.ajol.info//index.php/ijbcs/article/view/88422> |
| Weaverbirds | *Ploceus nelicourvi* | Incubation & nestling | Rakotomanana & Nakamura (2012), Breeding ecology of the Malagasy endemic Nelicourvi weaver *Ploceus nelicourvi*, *Ornithological Science,* 11: 39-46, <https://www.jstage.jst.go.jp/article/osj/11/1/11_39/_pdf/-char/ja> |
| Weaverbirds | *Ploceus taeniopterus* | Incubation | Jackson (1993), Causes of couspecific nest parasitism in the northern masked weaver. *Behavioral Ecology and Sociobiology,* 32**:** 119–126, <https://doi.org/10.1007/BF00164044> |
| Icterids | *Agelaius assimilis* | Incubation & nestling | Cooney et al. (2020). Ecology and allometry predict the evolution of avian developmental durations. *Nature Communications*, 11 (2383), <http://doi.org/10.1038/s41467-020-16257-x> |
| Icterids | *Amblycercus holosericeus* | Incubation & nestling | Cooney et al. (2020). Ecology and allometry predict the evolution of avian developmental durations. *Nature Communications*, 11 (2383), <http://doi.org/10.1038/s41467-020-16257-x> |
| Icterids | *Dolichonyx oryzivorus* | Nestling | Cooney et al. (2020). Ecology and allometry predict the evolution of avian developmental durations. *Nature Communications*, 11 (2383), <http://doi.org/10.1038/s41467-020-16257-x> |
| Icterids | *Icterus cucullatus* | Incubation | Magalhaes et al. (2009). A database of vertebrate longevity records and their relation to other life-history traits. *Journal of Evolutionary Biology*, *22*: 1770–1774. <http://doi.org/10.1111/j.1420-9101.2009.01783.x> |
| Icterids | *Icterus gularis* | Nestling | Werner et al. (2007), Breeding ecology of the Altamira oriole in the lower Rio Grande valley, Texas, *The Condor*, 109: 907-919, <https://doi.org/10.1093/condor/109.4.907> |
| Icterids | *Icterus icterus* | Incubation | Pujol & Mermoz (2011). Do life-history traits in the ancestor of cowbirds (*Molothrus* spp.) predispose them to become brood parasites? *Ornitologia Neotropical*, 22: 553-568. <https://sora.unm.edu/node/133265> |
| Icterids | *Quiscalus major* | Nestling | Cooney et al. (2020). Ecology and allometry predict the evolution of avian developmental durations. *Nature Communications*, 11 (2383), <http://doi.org/10.1038/s41467-020-16257-x> |

*Details and additional sources of life history data obtained from targeted literature searches.*

**SI Table 2**

| **Dep. variable** | **Ind. variable** | **Estimate** | **S.E.** | **T-value** | **p-value** | **N** | **R^2^** | **λ** |
| --- | --- | --- | --- | --- | --- | --- | --- | --- |
| Dev. period | Nest height | 0.02 | 0.02 | 1.03 | 0.31 | 51 | 0.16 | 0.00 |
|  | Body mass | 0.10 | 0.04 | 2.56 | 0.01 |  |  |  |
| Incubation period | Nest height | 0.01 | 0.01 | 1.05 | 0.30 | 52 | 0.06 | 0.00 |
|  | Body mass | 0.04 | 0.03 | 1.23 | 0.22 |  |  |  |
| Nestling period | Nest height | 0.02 | 0.02 | 0.89 | 0.38 | 52 | 0.14 | 0.00 |
|  | Body mass | 0.14 | 0.06 | 2.36 | 0.02 |  |  |  |

*Full results from models predicting log10 developmental period length, log10 incubation period length and log10 nestling period length from log10 nest height off the ground (or water) and log10 body mass in weaverbirds.*

**SI Table 3**

| **Dep. variable** | **Ind. variable** | **Estimate** | **S.E.** | **T-value** | **p-value** | **N** | **R^2^** | **λ** |
| --- | --- | --- | --- | --- | --- | --- | --- | --- |
| Dev. period | Colonial nesting | -0.01 | 0.01 | -0.96 | 0.34 | 56 | 0.10 | 0.38 |
|  | Body mass | 0.09 | 0.04 | 2.11 | 0.04 |  |  |  |
| Incubation period | Colonial nesting | -0.01 | 0.01 | -0.71 | 0.48 | 58 | 0.05 | 0.00 |
|  | Body mass | 0.05 | 0.03 | 1.58 | 0.11 |  |  |  |
| Nestling period | Colonial nesting | -0.02 | 0.02 | -0.95 | 0.35 | 57 | 0.06 | 0.51 |
|  | Body mass | 0.10 | 0.06 | 1.57 | 0.12 |  |  |  |

*Full results from models predicting log10 developmental period length, log10 incubation period length and log10 nestling period length from coloniality and log10 body mass in weaverbirds.*

**SI Table 4**

| **Dep. variable** | **Ind. variable** | **Estimate** | **S.E.** | **T-value** | **p-value** | **N** | **R^2^** | **λ** |
| --- | --- | --- | --- | --- | --- | --- | --- | --- |
| Dev. period | Nesting in thorns | 0.01 | 0.01 | 0.64 | 0.53 | 56 | 0.14 | 0.00 |
|  | Body mass | 0.10 | 0.04 | 2.61 | 0.01 |  |  |  |
| Incubation period | Nesting in thorns | -0.01 | 0.01 | -1.00 | 0.32 | 58 | 0.06 | 0.00 |
|  | Body mass | 0.06 | 0.03 | 1.78 | 0.08 |  |  |  |
| Nestling period | Nesting in thorns | 0.03 | 0.02 | 1.48 | 0.14 | 57 | 0.15 | 0.00 |
|  | Body mass | 0.12 | 0.06 | 2.16 | 0.04 |  |  |  |

*Full results from models predicting log10 developmental period length, log10 incubation period length and log10 nestling period length from nesting in thorny vegetation and log10 body mass in weaverbirds.*

**SI Table 5**

| **Dep. variable** | **Ind. variable** | **Estimate** | **S.E.** | **T-value** | **p-value** | **N** | **R^2^** | **λ** |
| --- | --- | --- | --- | --- | --- | --- | --- | --- |
| Dev. period | Nesting over water | 0.01 | 0.01 | 0.84 | 0.40 | 56 | 0.15 | 0.00 |
|  | Body mass | 0.10 | 0.04 | 2.82 | 0.01 |  |  |  |
| Incubation period | Nesting over water | 0.02 | 0.01 | 1.96 | 0.05 | 58 | 0.11 | 0.00 |
|  | Body mass | 0.05 | 0.03 | 1.53 | 0.13 |  |  |  |
| Nestling period | Nesting over water | 0.01 | 0.02 | 0.42 | 0.68 | 57 | 0.06 | 0.34 |
|  | Body mass | 0.11 | 0.06 | 1.82 | 0.07 |  |  |  |

*Full results from models predicting log10 developmental period length, log10 incubation period length and log10 nestling period length from nesting over water and log10 body mass in weaverbirds.*

**SI Table 6**

| **Dep. variable** | **Ind. variable** | **Estimate** | **S.E.** | **T-value** | **p-value** | **N** | **R^2^** | **λ** |
| --- | --- | --- | --- | --- | --- | --- | --- | --- |
| Incubation period | Nesting over water | 0.01 | 0.01 | 1.45 | 0.15 | 58 | 0.15 | 0.00 |
|  | Tunnel | 0.02 | 0.01 | 1.68 | 0.10 |  |  |  |
|  | Body mass | 0.04 | 0.03 | 1.31 | 0.20 |  |  |  |

*Full results from models predicting log10 developmental period length, log10 incubation period length and log10 nestling period length from nesting over water, nest tunnels and log10 body mass in weaverbirds.*

**SI Table 7**

| **Dep. variable** | **Ind. variable** | **Estimate** | **S.E.** | **T-value** | **p-value** | **N** | **R^2^** | **λ** |
| --- | --- | --- | --- | --- | --- | --- | --- | --- |
| Dev. period | Stinging insects | 0.02 | 0.01 | 1.37 | 0.18 | 56 | 0.17 | 0.00 |
|  | Body mass | 0.09 | 0.04 | 2.44 | 0.02 |  |  |  |
| Incubation period | Stinging insects | 0.00 | 0.01 | 0.23 | 0.82 | 58 | 0.04 | 0.00 |
|  | Body mass | 0.05 | 0.03 | 1.44 | 0.16 |  |  |  |
| Nestling period | Stinging insects | 0.03 | 0.02 | 1.39 | 0.17 | 57 | 0.14 | 0.00 |
|  | Body mass | 0.12 | 0.06 | 2.18 | 0.03 |  |  |  |

*Full results from models predicting log10 developmental period length, log10 incubation period length and log10 nestling period length from nesting in association with stinging insects and log10 body mass in weaverbirds.*

**SI Table 8**

| **Dep. variable** | **Ind. variable** | **Estimate** | **S.E.** | **T-value** | **p-value** | **N** | **R^2^** | **λ** |
| --- | --- | --- | --- | --- | --- | --- | --- | --- |
| Dev. period | Nest height | 0.05 | 0.02 | 2.17 | 0.04 | 34 | 0.43 | 0.74 |
|  | Body mass | 0.16 | 0.04 | 3.93 | <0.01 |  |  |  |
| Incubation period | Nest height | 0.04 | 0.02 | 2.43 | 0.02 | 36 | 0.45 | 0.94 |
|  | Body mass | 0.12 | 0.03 | 4.27 | <0.01 |  |  |  |
| Nestling period | Nest height | 0.13 | 0.03 | 4.08 | <0.01 | 38 | 0.40 | 0.00 |
|  | Body mass | 0.17 | 0.06 | 2.83 | <0.01 |  |  |  |

*Full results from models predicting log10 developmental period length, log10 incubation period length and log10 nestling period length from log10 nest height off the ground (or water) and log10 body mass in icterids.*

**SI Table 9**

| **Dep. variable** | **Ind. variable** | **Estimate** | **S.E.** | **T-value** | **p-value** | **N** | **R^2^** | **λ** |
| --- | --- | --- | --- | --- | --- | --- | --- | --- |
| Dev. period | Nest height | 0.02 | 0.02 | 1.19 | 0.24 | 34 | 0.72 | 0.00 |
|  | Suspended | 0.04 | 0.02 | 2.30 | 0.03 |  |  |  |
|  | Pendulous | 0.14 | 0.02 | 5.54 | <0.01 |  |  |  |
|  | Body mass | 0.13 | 0.03 | 3.76 | <0.01 |  |  |  |
| Incubation period | Nest height | 0.03 | 0.02 | 2.11 | 0.04 | 36 | 0.58 | 0.97 |
|  | Suspended | 0.01 | 0.02 | 0.35 | 0.73 |  |  |  |
|  | Pendulous | 0.06 | 0.03 | 2.42 | 0.02 |  |  |  |
|  | Body mass | 0.10 | 0.03 | 3.62 | <0.01 |  |  |  |
| Nestling period | Nest height | 0.10 | 0.03 | 3.06 | <0.01 | 38 | 0.57 | 0.00 |
|  | Suspended | 0.03 | 0.03 | 0.92 | 0.37 |  |  |  |
|  | Pendulous | 0.16 | 0.05 | 3.47 | <0.01 |  |  |  |
|  | Body mass | 0.13 | 0.06 | 2.08 | 0.05 |  |  |  |

*Full results from models predicting log10 developmental period length, log10 incubation period length and log10 nestling period length from log10 nest height off the ground (or water), nest type and log10 body mass in icterids.*

**SI Table 10**

| **Dep. variable** | **Ind. variable** | **Estimate** | **S.E.** | **T-value** | **p-value** | **N** | **R^2^** | **λ** |
| --- | --- | --- | --- | --- | --- | --- | --- | --- |
| Dev. period | Colonial nesting | 0.00 | 0.02 | 0.16 | 0.87 | 48 | 0.29 | 0.85 |
|  | Body mass | 0.18 | 0.04 | 4.26 | <0.01 |  |  |  |
| Incubation period | Colonial nesting | -0.01 | 0.01 | -0.79 | 0.43 | 50 | 0.26 | 0.75 |
|  | Body mass | 0.11 | 0.03 | 4.01 | <0.01 |  |  |  |
| Nestling period | Colonial nesting | 0.02 | 0.03 | 0.57 | 0.57 | 52 | 0.14 | 0.70 |
|  | Body mass | 0.19 | 0.07 | 2.68 | 0.01 |  |  |  |

*Full results from models predicting log10 developmental period length, log10 incubation period length and log10 nestling period length from coloniality and log10 body mass in icterids.*

**SI Table 11**

| **Dep. variable** | **Ind. variable** | **Estimate** | **S.E.** | **T-value** | **p-value** | **N** | **R^2^** | **λ** |
| --- | --- | --- | --- | --- | --- | --- | --- | --- |
| Dev. period | Nesting in thorns | 0.01 | 0.01 | 0.40 | 0.69 | 48 | 0.29 | 0.85 |
|  | Body mass | 0.17 | 0.04 | 3.99 | <0.01 |  |  |  |
| Incubation period | Nesting in thorns | 0.02 | 0.01 | 1.53 | 0.13 | 50 | 0.28 | 0.78 |
|  | Body mass | 0.10 | 0.03 | 3.42 | <0.01 |  |  |  |
| Nestling period | Nesting in thorns | -0.01 | 0.03 | -0.47 | 0.64 | 52 | 0.13 | 0.73 |
|  | Body mass | 0.20 | 0.08 | 2.65 | 0.01 |  |  |  |

*Full results from models predicting log10 developmental period length, log10 incubation period length and log10 nestling period length from nesting in thorny vegetation and log10 body mass in weaverbirds.*

**SI Table 12**

| **Dep. variable** | **Ind. variable** | **Estimate** | **S.E.** | **T-value** | **p-value** | **N** | **R^2^** | **λ** |
| --- | --- | --- | --- | --- | --- | --- | --- | --- |
| Dev. period | Nesting over water | -0.02 | 0.02 | -1.13 | 0.27 | 48 | 0.31 | 0.85 |
|  | Body mass | 0.17 | 0.04 | 4.22 | <0.01 |  |  |  |
| Incubation period | Nesting over water | -0.01 | 0.01 | -1.06 | 0.29 | 50 | 0.27 | 0.70 |
|  | Body mass | 0.11 | 0.03 | 3.93 | <0.01 |  |  |  |
| Nestling period | Nesting over water | -0.04 | 0.03 | -1.62 | 0.11 | 52 | 0.16 | 0.75 |
|  | Body mass | 0.18 | 0.07 | 2.49 | 0.02 |  |  |  |

*Full results from models predicting log10 developmental period length, log10 incubation period length and log10 nestling period length from nesting over water and log10 body mass in icterids.*
